# Cytotoxic activity against human leukemia cells of Kaempferol-3-O-rhamnoside from Vietnamese *Schima wallichii* (DC.) Korth: A combination of *in vitro* and *in silico* insights

**DOI:** 10.1101/2024.12.25.630323

**Authors:** Manh Hung Tran, Yen Nhi Nguyen, Thuy Mi Pham Lam, Thuy Linh Thi Tran, Tan Khanh Nguyen, Tuan Anh Le, Van Ngo Thai Bich, Hieu Phu Chi Truong, Phu Tran Vinh Pham

## Abstract

In the investigation of the cytotoxic activity against leukemia cells of Vietnamese medicinal plants, we identified the extract of *Schima wallichii* (DC.) Korth as capable of inhibiting several leukemia cell lines. In this study, we isolated a main compound as kaempferol-3-O-rhamnoside, marking the first report of this compound being isolated from the stem of *Schima wallichii* collected in Vietnam. In *in vitro* experiments, kaempferol-3-O-rhamnoside exhibited cytotoxic effects on three leukemia cell lines, HL-60 and KG-1. Regarding its mechanism of action, the compound effectively inhibited growth of HL-60 and KG-1 leukemia cell lines by activating caspase-3 and caspase-9 in both cell lines. Additionally, kaempferol-3-O-rhamnoside upregulated the pro-apoptotic protein Bax while downregulating the anti-apoptotic protein Bcl-2 in these cell lines. *In silico* experiments revealed that docking simulations showed kaempferol-3-O-rhamnoside binds to both the allosteric site of procaspase-3 and the active site of PARP1, with binding energies of -7.36 and -10.76 kcal/mol, respectively. Kaempferol-3-O-rhamnoside demonstrated stable binding affinity with PARP1, characterized by significant hydrogen bonding, hydrophobic interactions, and pi-stacking in the molecular dynamic simulations. These results suggest that kaempferol-3-O-rhamnoside has the potential PARP1 inhibitor, making it a promising candidate for targeting leukemia cells. Moreover, it provides evidence for considering this compound in drug discovery and development targeting PARP1-related pathways.

## Introduction

Leukemia is a type of blood cancer characterized by abnormal white blood cells in the bone marrow, rapidly invading healthy cells through the bloodstream [1]. Most cases of leukemia are acute and fast-growing, while some are chronic and progress more slowly. Acute leukemia accounts for approximately 75% of cases and is a malignant disorder affecting lymphocytes in both children and adults [2,3]. Apoptosis, or programmed cell death, is a natural process that plays a crucial role in cell replacement, tissue regeneration, and removing damaged cells under normal conditions [4]. Many studies and applications have aimed to enhance or reduce cell sensitivity to apoptosis, forming the basis for treatments for various diseases [5]. However, dysregulation of apoptosis can lead to severe disorders such as neurodegenerative diseases, autoimmune diseases, and cancer [6,7]. Research indicates that most cancer cases result from abnormal cell genetic changes during development [8,9]. Recent studies have highlighted that leukemia cells consistently exhibit abnormalities in one or more apoptotic pathways, providing these cells with a survival advantage over normal cells [10]. Development leukemia deregulation of apoptosis disrupts the intricate balance between cell proliferation, survival, and death, playing a crucial role in the development of diseases such as cancer, particularly leukemia, which is recognized as a multistep process, marked by a series of genetic alterations [11]. The disease originates from a single cell that undergoes malignant transformation due to frequent genetic mutations, followed by the clonal selection of mutated cells, which increasingly exhibit aggressive behavior [12,13].

Apoptosis can be triggered by several factors, including the activation of the caspase enzyme system [14]. The timing of caspase activation and mitochondrial remodeling by cytochrome C can differ depending on the mechanism [15]. One crucial mechanism identified by scientists is the activation of caspase enzymes before mitochondrial changes occur. The oligomerization of receptors leads to the activation of caspase-8, which subsequently activates caspase-3, resulting in cell death [14]. Other cancer cell death pathways are triggered by both extracellular and intracellular factors, such as growth factors, hypoxia, DNA damage, and oncogene activation. Studies on apoptotic cell death have consistently shown that the activation of caspase-9, followed by caspase-3, induces apoptosis [14,15]. Caspases can also cleave various cellular structures, including cytoskeletal components, nuclear proteins, and precursor proteins such as lamin, actin, and ADP-ribose-binding protein (PARP) [14]. The cleavage of these proteins is associated with morphological changes in cells, which have been well-documented during apoptosis [16].

Poly ADP-ribose polymerase 1 (PARP1) is a nuclear enzyme crucial for regulating several cellular processes, such as DNA repair, transcription, and chromatin remodeling. It is a member of the ADP-ribosyltransferase enzyme family, which modifies proteins by attaching ADP-ribose units [17]. In cancer therapy, inhibiting PARP1 hampers the ability of cancer cells to repair DNA damage, leading to cell death. Normal cells, however, are better equipped to tolerate PARP1 inhibition due to functional DNA repair mechanisms. Several PARP1 inhibitors, such as olaparib, rucaparib, niraparib, and talazoparib, have been approved for treating various cancers, particularly ovarian and breast cancer, and have demonstrated promising outcomes in clinical trials [17–19].

Although modern medicine has made significant advancements in the treatment of leukemia, disease recurrence and drug resistance remain major challenges, necessitating more effective treatment approaches. Currently, most anti-cancer agents induce programmed cell death (apoptosis). However, while these agents can effectively kill cancer cells, they may also be toxic to normal cells. This highlights the urgent need to discover new compounds that can treat cancer with minimal side effects and help prevent drug resistance. Although modern medical treatments, such as radiotherapy, chemotherapy, immunotherapy, and bone marrow or stem cell transplantation, are widely used for cancer management, combining these approaches with natural herbs and traditional remedies can enhance patient resilience, reduce the toxicity associated with chemotherapy and radiotherapy, and promote faster recovery [19,20]. Scientific research has shown that natural products possess the potential to treat or support the treatment of cancer effectively [21–24].

Our screening process revealed that the extract of *Schima wallichii* (DC.) Korth, a member of the Theaceae family found in Quang Tri, has potential inhibitory effects on the growth and development of leukemia cell lines. This led to the hypothesis that compounds in *Schima wallichii* and serrated myrtle may possess *in vitro* anti-leukemia activity. *Schima wallichii*, also known as Khao cai or Thu lu, has the scientific name *Schima wallichii* (DC.) Korth (synonym: *Gordonia wallichii* DC.) belongs to the *Schima* genus within the Theaceae family [25]. These are large trees, reaching heights of 20-25 m, and are widely distributed in Vietnam as well as in other countries such as Brunei, China, India, Indonesia, Laos, Malaysia, Myanmar, Nepal, Papua New Guinea, the Philippines, and Thailand [26,27]. In Vietnam, *Schima wallichii* is commonly found in the highland forests of provinces like Lao Cai, Lai Chau, Cao Bang, and several areas in the Central Highlands. Trees of the *Vối* genus are notable for their large size and durable wood, which is classified as Group V [28,29]. The wood is known for being heavy, resistant to warping and termites, and has an attractive brown color, making it useful for building house pillars and household items. The bark, leaves, and roots are traditionally used to treat various ailments and for producing industrial products. *Schima wallichii* (Vối Thuốc) is also cultivated as an effective green fireproof barrier. It is a sun-loving tree, grows relatively quickly, adapts to various terrains, and has strong natural regeneration capabilities. Due to these advantageous characteristics, *Schima wallichii* is among the few tree species prioritized for upstream protective forest planting. In traditional medicine, the bark and young leaves of *Schima wallichii* are commonly used for medicinal purposes. The bark serves as a wound disinfectant, anthelmintic, and treatment for gonorrhea [28,29]. A decoction of the bark is known to reduce fever and prevent lice infestations [29]. The young leaves, which have an astringent taste, neutral properties, and mild toxicity, are used to kill roundworms, act as antiseptics and antifungals, and treat headaches while supporting cancer treatment. The flowers, seeds, and roots have demonstrated antioxidant, antibacterial, anti-inflammatory, and analgesic properties. Literature reviews have shown that *Schima wallichii* possesses potent antioxidant activity [30]. To date, extracts and certain compounds from *Schima wallichii* have been evaluated for their significant biological effects, including antibacterial, antiviral, antimalarial, and anticancer properties [25,31–33].

Currently, there are no comprehensive scientific studies focused on the chemical composition of this plant extraction and investigation of the mechanisms by which *Schima wallichii* inhibits leukemia cell lines, especially via the apoptosis pathway. Therefore, building on existing literature and prior scientific findings, this article presents research on identifying and discovering additional natural compounds from this plant species and then predicting the action mechanism of these compounds on leukemia cells based on *in vitro* and *in silico* investigations. The goal is to uncover new active ingredients with potential anti-leukemia properties, laying the groundwork for specialized pharmaceutical research and developing these active compounds as supportive agents or drugs for leukemia treatment.

## Materials and Methods

### Plant materials

*Schima wallichii* (DC.) Korth was collected from the mountainous region of Lao Bao District, Quang Tri Province, Vietnam in October 2020. The plant specimens were identified by Dr. Tuan Anh Le of the Vietnam Academy of Science and Technology (VAST). The collected samples are currently stored in the Laboratory of the Department of Pharmacognosy and Drug Control at the School of Medicine and Pharmacy, The University of Danang, Vietnam.

### Extraction and isolation

The plant branches were dried and cut into small pieces (2.0 kg) before being refluxed with 5 liters of 70% ethanol in a 10-liter flask. This process was repeated three times, each lasting 3 hours. The resulting extract was filtered and evaporated under reduced pressure, yielding a total of 118.6 g of extract. The entire extract was dissolved in 1 liter of warm water and partitioned with n-hexane, CH_2_Cl_2_, and EtOAc. Each phase was evaporated under reduced pressure, resulting in the following fractions: n-hexane (2.15 g), CH_2_Cl_2_ (10.54 g), EtOAc (28.50 g), and an aqueous residue (W). The EtOAc extract (28.50 g) was further separated on a normal-phase silica gel column and eluted with a CH_2_Cl_2_-MeOH solvent system at different ratios (50:1, 25:1, 10:1, 5:1, 1:1, 0:1). Each stage used 1.0 liter of the mixture, resulting in 10 smaller fractions (E1 - E10). Sub-fraction E8 (318 mg) was subjected to normal-phase column chromatography (column size 1.5 x 60 cm) with a mobile phase of CH_2_Cl_2_-Ac (5:1 to 1:1) to isolate compound 4 (28.5 mg). Detailed NMR spectra for each isolated compound are provided in the S1_File. Kaempferol-3-O-rhamnoside: Light yellow powder, soluble in water, methanol and ethanol, melting point 216— 218°C; IR (KBr)max: 3512, 3490, 2960, 2881, 1750, 1148 cm^-1^; FAB-MS [M]^+^ 416.1 *m*/*z* (C_21_H_20_O_9_). ^1^H-NMR (MeOD, 400 MHz) δ_H_: 6.17 (1H, d, *J*=2.4 Hz, H-6), 6.34 (1H, d, *J*=2.4 Hz, H-8), 7.73 (2H, dd, *J*=1.8, 8.0 Hz, H-2’, 6’), 6.92 (2H, dd, *J*=1.8, 8.0 Hz, H-3’, 5’), Rha: 4.23 (1H, d, *J*=2.5 Hz, H-1”), 3.31-3.39 (m, H-2”, 3”, 4”, 5”), 0.92 (3H, d, *J*=5.7 Hz, H-6”). ^13^C-NMR (MeOD, 100 MHz) δ_C_: 163.2 (C-2), 159.3 (C-3), 179.6 (C-4), 161.6 (C-5), 103.5 (C-6), 163.2 (C-7), 94.8 (C-8), 156.5 (C-9), 106.0 (C-10), 122.7 (C-1’), 132.0 (C-2’, C-6’), 116.3 (C-3’, 5’), 163.2 (C-4’), Rha: 103.5 (C-1’’), 72.4 (C-2’’), 73.3 (C-3’’),73.7 (C-4’’), 72.0 (C-5’’), 17.7 (C-6’’).

### Cancer cell lines

The human acute myeloid leukemia cell line HL-60, cervical epithelial cell line HeLa, human embryonic kidney cell line HEK-293, and the human acute myeloid leukemia cell line KG-1 were obtained from the American Type Culture Collection (ATCC, Manassas, VA, USA). HL-60 cells were cultured in Roswell Park Memorial Institute (RPMI, Gibco-USA) 1640 medium supplemented with 10% fetal bovine serum (FBS, Gibco-USA), 100 U/mL penicillin, and 100 µg/mL streptomycin. The cultures were maintained at 37°C in a 5% CO2 incubator. Cells were subcultured by diluting the cell suspension in fresh medium to a density of 2 × 10^5^ cells/mL. HL-60 leukemia cells are non-adherent and grow in suspension in the culture medium. Subculturing was performed 2-3 times per week when the cell density reached 80% of the culture volume.

### Antiproliferative assays

The antiproliferative activity of kaempferol-3-O-rhamnoside was evaluated in human acute myeloid leukemia cell lines HL-60 and KG-1, as well as other cancer cell lines, including pancreatic (MIA PaCa-2, PANC-1), lung (A549), ovarian (OVCAR-8), and cervical cancer (HeLa), using an improved Dojindo kit based on the conventional MTT assay. Cells were cultured in 96-well plates at 37°C in a 5% CO2 environment. Kaempferol 3-O-rhamnoside samples were diluted in dimethyl sulfoxide (DMSO, Sigma-USA) to various concentrations, with the final DMSO concentration kept below 0.1% to prevent cell differentiation. After 24 hours of cell incubation, Kaempferol 3-O-rhamnoside was added to each well, and the same medium with 0.1% DMSO was added to the blank control wells (without a test sample). After 48 hours of incubation, the Dojindo reagent was added to each well. Four hours later, the optical density (OD) was measured at 450 nm using a microplate reader (Molecular Devices, USA). The assays were conducted in triplicate to ensure the accuracy and reliability of the results. Camptothecin (Merck) was used as a positive control, while DMSO served as a negative control. IC50 values (the concentration required to inhibit 50% of cell growth) were calculated using TableCurve 2Dv5.01 software [34].

### Caspase-3 activation

The experiment was performed following the protocols previously described by Vo et al. (2021) [34]. Briefly, HL-60 cancer cells were treated with kaempferol-3-O-rhamnoside at concentrations of 3, 10, and 30 μM. After 12, 24, and 48 hours of incubation, the cells were collected, washed with cold Phosphate-buffered saline (PBS, Gibco-USA), and lysed with lysis buffer at room temperature for 10 minutes. The lysates were then cooled and mixed with assay buffer in each well and incubated at 37°C for 1 hour. Fluorescence intensity was measured using a Twinkle LB970 spectrophotometer (Berthold Technologies, Germany) at an excitation wavelength of 370 nm and an emission wavelength of 505 nm. The control group consisted of untreated cancer cells. The level of caspase-3 activation was determined by comparing the fluorescence intensity of the treated samples to that of the control. Data analysis was performed using GraphPad Prism 3.03 software.

### Western Blotting assay

HL-60 and KG-1 cells (10^6^ cells/well) were treated with kaempferol 3-O-rhamnoside at concentrations of 3, 10, and 30 µM for 24 hours at 37°C. The cell lysates were prepared in a lysis buffer containing a protease inhibitor cocktail (Roche, Germany) [34]. The cell debris was removed by centrifugation at 14,000 rpm for 10 minutes. The protein concentration in the supernatant was measured using the Bio-Rad DC Protein Assay kit (Bio-Rad, USA). The protein extracts were separated by Sodium Dodecyl Sulphate-Polyacrylamide Gel Electrophoresis (SDS-PAGE) and transferred onto a Polyvinylidene fluoride (PVDF) membrane (Bio-Rad, USA). The membrane was blocked with 5% nonfat dry milk for 1 hour and washed three times with TBS-T (Tris-buffered saline containing 0.1% Tween-20) at room temperature for 10 minutes each. The membrane was then incubated with primary antibodies targeting Bax, Bcl-2, Cytochrome-c, activated caspase-3, activated caspase-9, and cleaved PARP1 (Cell Signaling Technology, USA) at 4°C overnight. After washing three times with TBS-T, the membrane was incubated with HRP (horseradish peroxidase)-conjugated secondary antibodies at room temperature for 1.5 hours, followed by three additional washes with TBS-T. Protein bands were visualized using an enhanced chemiluminescence (ECL) kit as per the manufacturer’s instructions (Santa Cruz Biotechnology) and detected by X-ray exposure. Anti-β-actin antibody was used as a loading control. The used membrane was soaked in membrane washing buffer (Gene Bio-Application Ltd.) at room temperature for 20 minutes.

### Protein and ligand preparation

The crystal structure of PARP1 (PDB ID: 5WS1) was retrieved from the Protein Data Bank (RCSB PDB). Extraneous water molecules and co-crystallized ligands were removed using Discovery Studio 2020 Client. Hydrogen atoms were added to the protein using AutoDockTools 1.5.6, and the protein structures were exported in pdbqt format for docking. The 3D structures of the compounds were obtained from the PubChem library, with hydrogen atoms added and conversion to pdbqt format performed using Open Babel software for screening.

### Docking simulation

The structure files of the compounds were generated using MarvinSketch (ChemAxon, Budapest, Hungary). Protein structures were edited using UCSF Chimera 1.14 to remove water molecules, rebuild residues, and eliminate co-crystallized ligands and cofactors. The proteins and compounds were then prepared using AutoDockTools 1.5.6. Ligand binding sites and grid box parameters were determined based on potential binding cavities identified by MetaPocket2.0. Molecular docking was performed using AutoDock 4.2 with the Lamarckian genetic algorithm and default settings. The docking results were analyzed with Discovery Studio Visualizer.

### Molecular dynamics simulations

Molecular dynamics (MD) simulations were conducted using GROMACS 2020.4 to evaluate the stability of the ligand-protein complex [35]. The bound complex was prepared using the CHARMM36 force field and positioned in an octahedral box filled with TIP3P water molecules. K+ and Cl-ions were added to neutralize the system. Energy minimization was performed using the Steepest Descent Method with a maximum force criterion of < 1000 kJ/mol/nm over 5000 steps. The systems were equilibrated under isothermal-isochoric (NVT) and isothermal-isobaric (NPT) conditions for 100 ps and 1000 ps, respectively, ensuring stability at 300 K and 1 bar. Finally, 100 ns MD simulations were carried out for each complex.

### Binding free energy estimation

The MM/PBSA (Molecular Mechanics/Poisson–Boltzmann Surface Area) method was used to estimate the relative binding free energy of the compound and assess the stability of the apo-protein. In this study, MM/PBSA calculations were performed using the MOLAICAL script to determine the relative binding free energy [36].

### Statistical analysis

Analysis of variance (ANOVA) followed by Tukey’s test was performed on pre-validated data. Additionally, statistical analysis was conducted using Prism (GraphPad Software, San Diego, CA, USA) with a t-test for comparisons between two groups or ANOVA followed by Bonferroni post-hoc analysis for multiple group comparisons and correlation analysis. Data were expressed as mean ± standard deviation (SD). Statistical significance was considered at *** p < 0.001; ** p < 0.05; * p < 0.1.

## Results and Discussion

### Kaempferol-3-O-rhamnoside determination

This compound was obtained as a pale yellow powder, highly soluble in water, methanol, and ethanol, with a melting point of 216—218 °C. The IR spectrum (KBr) showed key absorption bands at 3512, 3490, 2960, 2881, 1750, and 1148 cm⁻¹, indicating the presence of hydroxy and carbonyl groups. Positive FAB-MS analysis revealed a molecular ion peak at [M]+ 416.1 m/z, consistent with the molecular formula C_21_H_20_O_9_. ^1^H-NMR (MeOD, 400 MHz). The spectrum displayed signals characteristic of the flavonoid skeleton, including δH 6.17 (1H, d, J = 2.4 Hz, H-6), 6.34 (1H, d, J = 2.4 Hz, H-8), and aromatic signals at δH 7.73 (2H, dd, J = 1.8, 8.0 Hz, H-2’, 6’) and 6.92 (2H, dd, J = 1.8, 8.0 Hz, H-3’, 5’). Signals corresponding to the rhamnose sugar moiety were observed at δH 4.23 (1H, d, J = 2.5 Hz, H-1’’) and 3.31-3.39 (m, H-2’’, 3’’, 4’’, 5’’), with a notable doublet at δH 0.92 (3H, d, J = 5.7 Hz, H-6’’), confirming the rhamnose structure. The ^13^C spectrum (MeOD, 100 MHz) and DEPT analysis revealed a total of 21 carbon signals. Six signals were assigned to the rhamnose sugar group at δC 103.5 (C-1’’), 72.4 (C-2’’), 73.3 (C-3’’), 73.7 (C-4’’), 72.0 (C-5’’), and 17.7 (C-6’’). The remaining 15 signals were attributed to the aglycone backbone: δC 163.2 (C-2), 159.3 (C-3), 179.6 (C-4), 161.6 (C-5), 103.5 (C-6), 163.2 (C-7), 94.8 (C-8), 156.5 (C-9), 106.0 (C-10), 122.7 (C-1’), 132.0 (C-2’, C-6’), 116.3 (C-3’, 5’), and 163.2 (C-4’). The presence of a carboxyl group was indicated by the signal at δC 179.6 (C-4), confirming the flavonoid structure. The comparison of ^1^H- and ^13^C-NMR data with literature values indicated that this compound has a kaempferol backbone (aglycone) with a rhamnose sugar attached. The glycosidation site at C-3 was inferred from the observed downfield shifts of the carbon signals. Acid hydrolysis result yielded L-rhamnose which was confirmed by TLC analysis. The heteronuclear multiple bond connectivity (HMBC) spectra showed correlations between H-1’’ of rhamnose and C-3 of the kaempferol backbone, confirming the attachment site. Based on these data, this compound was identified as kaempferol 3-O-rhamnoside (Fig 1) [32,33].

**Fig 1.**
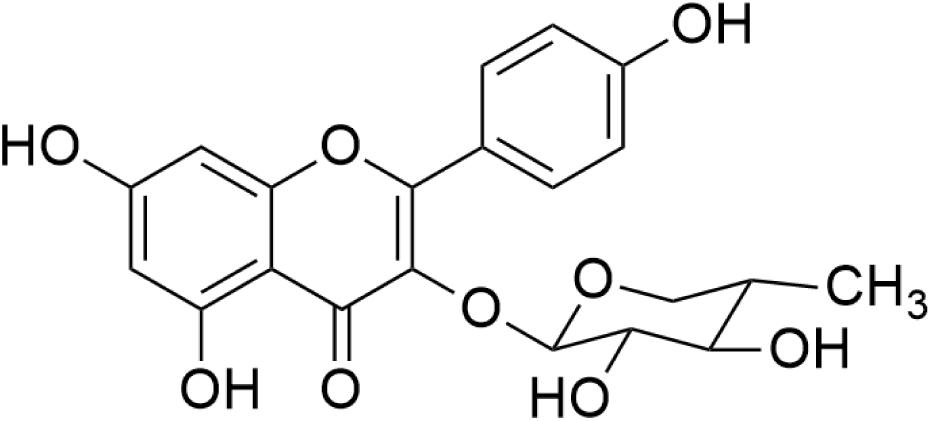
Structure of Kaempferol-3-O-rhamnoside from *Schima wallichii* Korth.

### Cytotoxic activity

The cytotoxicity of Kaempferol-3-O-rhamnoside was evaluated based on the activity against human leukemia cell lines HL-60 and KG-1, cervical cancer cell line HeLa, and human embryonic kidney cell line HEK-293 (Table 1).

**Table 1.** Cytotoxic activity of the testing compound.

| Compounds | Cancer cells, IC<sub>50</sub> μM<sup>a)</sup> |  |  |  |
| --- | --- | --- | --- | --- |
|  | HeLa | HL-60 | KG-1 | HEK-293 |
| Kaempferol-3-O-rhamnoside (4) | > 30 | 10,4 ± 1,0 | 11,5 ± 1,6 | > 30 |
| Camptothecin<sup>b)</sup> | 0.5 ± 0.1 | 1,7 ± 0,3 | 0,5 ± 0,1 | - |

<sup>a)</sup> IC<sub>50</sub> values were calculated based on the results of three independent assays to ensure repeatability and reliability. <sup>b)</sup> Camptothecin was used as the positive control. The data are presented as mean values ± standard deviation (S.D.). (-) indicates either weak activity or that the activity was not tested.

Among the tested cell lines, kaempferol-3-O-rhamnoside exhibited the strongest cytotoxic effects against KG-1 leukemia cell lines, with IC_50_ values of 11.5 ± 1.6 µM, followed by HL-60 with IC_50_ 10.4 ± 1.0 µM. However, the compound demonstrated weak cytotoxic activity against HeLa cells, with an IC_50_ value exceeding 30 μM. Due to the notable cytotoxicity of kaempferol-3-O-rhamnoside against leukemia cells, further research was conducted to investigate its molecular-level effects on these cells.

### Effect of Kaempferol-3-O-rhamnoside on the activation of caspase-3

At first, the effect of kaempferol-3-O-rhamnoside on the growth of leukemia cells was investigated. HL-60 cells were treated with varying concentrations of Kaempferol-3-O-rhamnoside (3–30 μM) for 48 hours. Cell viability was assessed through microscopy. The results indicated that exposure to kaempferol-3-O-rhamnoside for 24 hours significantly increased the number of cell deaths in HL-60 cells (Fig 2A).

**Fig 2.**
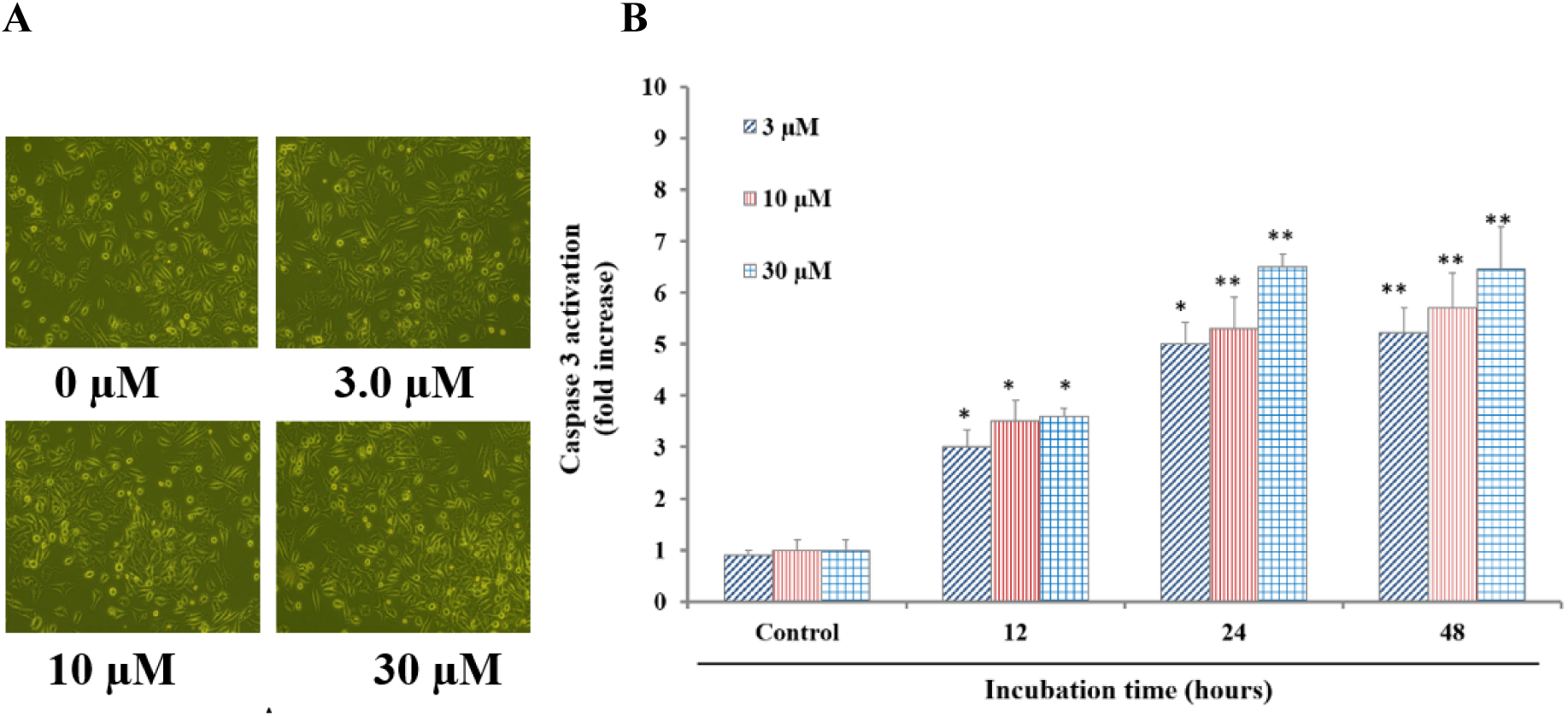
Effect of Kaempferol-3-O-rhamnoside (4) on leukemia cell death and caspase-3 activation. **A)** Viability of HL-60 cells. Cell morphology was visualized by microscopy (500 µm). Cells were seeded in 6-well plates at a density of 1 × 10^5^ cells/well and treated with the compound for 24 hours, **B)** Caspase-3 activation in HL-60 cells. The control group was treated with 0.1% DMSO. Data are presented as the mean ± standard deviation of three independent experiments (*p < 0.01; **p < 0.05).

Caspase-3 is a crucial enzyme in the apoptosis pathway, activated by both intrinsic and extrinsic signaling pathways. It is initially present as a 32 kDa precursor protein known as procaspase-3. Upon stimulation, caspase-3 becomes activated, leading to DNA fragmentation and apoptosis in cancer cells by cleaving DNA and proteins into smaller fragments. In this experiment, the substrate Ac-Asp-Glu-Val-Asp-8-amino-4-trifluoromethylcoumarin (Av-DEVD-AFC) was used at different time points (12, 24, and 48 hours) to evaluate caspase-3 activation in HL-60 cells (Fig 2B). After 12 hours of incubation, kaempferol-3-O-rhamnoside enhanced procaspase-3 activation, resulting in a 3-to 4-fold increase in cleaved caspase-3 levels compared to the control. After 24 hours, cleaved caspase-3 activation showed a 6-to 8-fold increase relative to the control. These findings suggest that kaempferol-3-O-rhamnoside promotes caspase-3 activation in a dose- and time-dependent manner, particularly during the early stages of exposure in HL-60 cells.

### Effect of kaempferol 3-O-rhamnoside on protein expression levels

To further evaluate the apoptotic effects of kaempferol-3-O-rhamnoside on HL-60 and KG-1 cells, the expression of cleaved poly(ADP-ribose) polymerase (PARP1) was examined after treatment with kaempferol-3-O-rhamnoside (0–30 μM) using Western blot analysis.

As shown in (Fig 3), kaempferol-3-O-rhamnoside induced the cleavage of PARP1, converting it into its cleaved form. This transformation inhibits PARP1’s ability to perform its DNA repair function, thereby preventing the restoration of DNA damage in cancer cells. Analysis of cleaved PARP1 content demonstrated that kaempferol-3-O-rhamnoside caused a concentration dependent increase in cleaved PARP1, leading to apoptosis in HL-60 and KG-1 cells during incubation.

**Fig 3.**
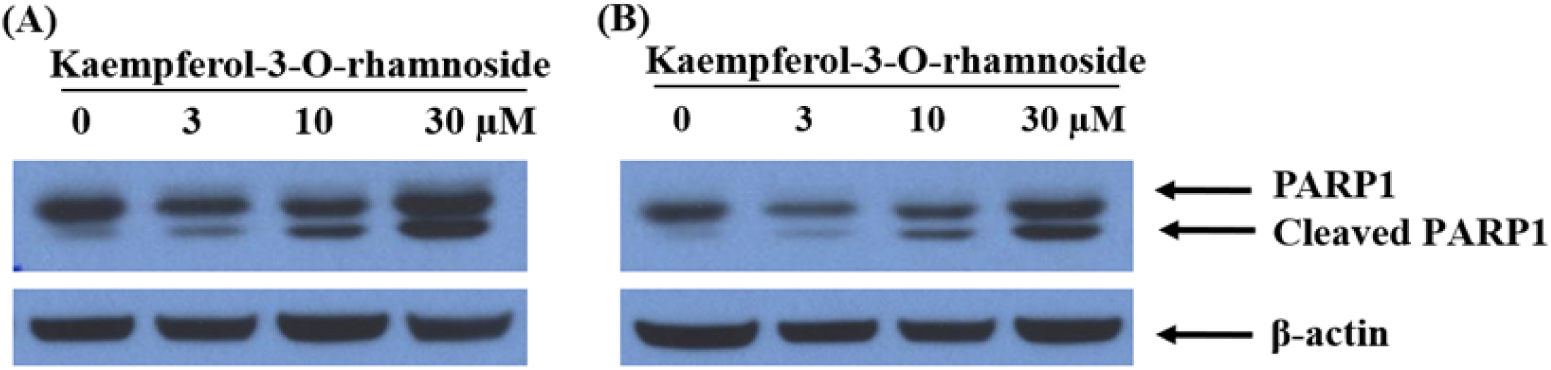
Effect of kaempferol 3-O–rhamnoside on PARP1 expression. **A)** HL-60 cells, **B)** KG-1 cells. Western blotting assays used anti-PARP1 and Cleaved PARP1 antibodies; β-Actin was used as an internal control.

To determine whether kaempferol-3-O-rhamnoside can induce apoptosis in HL-60 and KG-1 cells, the protein levels of caspase-3, caspase-9, and PARP1 were measured by Western blotting with specific antibodies (Fig 4). PARP proteins play a crucial role in DNA repair, genomic stability, and programmed cell death. Activation of caspase-3 often leads to PARP1 cleavage, which acts as a positive regulator of apoptosis and is considered a biomarker for apoptosis detection. HL-60 and KG-1 cells were treated with Kaempferol-3-O-rhamnoside (0–30 µM) for 48 hours. Proteins (50 µg per lane) from cell lysates were separated by SDS-PAGE and transferred to PVDF membranes. As shown in Fig. 4, kaempferol-3-O-rhamnoside induced the conversion of procaspase-3 (32 kDa) to active caspase-3 (19 kDa) and procaspase-9 (50 kDa) to active caspase-9 (37 kDa) in a dose-dependent manner. Additionally, after 48 hours of treatment, both HL-60 and KG-1 cells showed a gradual increase in cleaved PARP1 levels in a concentration-dependent manner. The results indicate that kaempferol-3-O-rhamnoside at concentrations as low as 3.0 µM can activate procaspase-3 and procaspase-9. With increasing concentrations (10–30 µM), the expression of active caspase-3 and caspase-9 further increased, suggesting that kaempferol-3-O-rhamnoside induces apoptosis in HL-60 and KG-1 cells through caspase activation.

**Fig 4.**
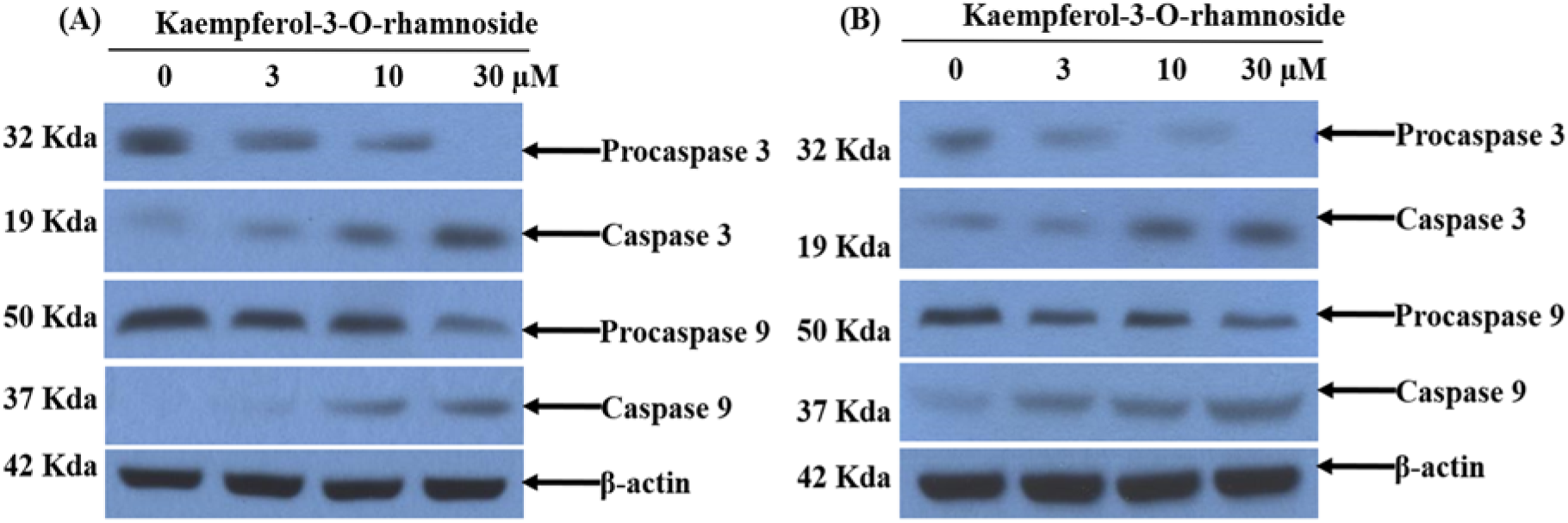
Effect of kaempferol 3-O–rhamnoside on caspase expressions. **A)** HL-60 cells, **B)** KG-1 cells. Western blotting assays used anti-Procaspase 3, Caspase 3, Procaspase 9, and Caspase 9 antibodies; β-Actin was used as an internal control

The effects of kaempferol 3-O-rhamnoside on the expression levels of pro-apoptotic protein Bax, anti-apoptotic protein Bcl-2, and mitochondrial damage-associated cytochrome c (Cyt-c)—which is released from mitochondria into the cytoplasm to activate the apoptotic process—were studied in HL-60 and KG-1 cancer cells (Fig 5).

**Fig 5.**
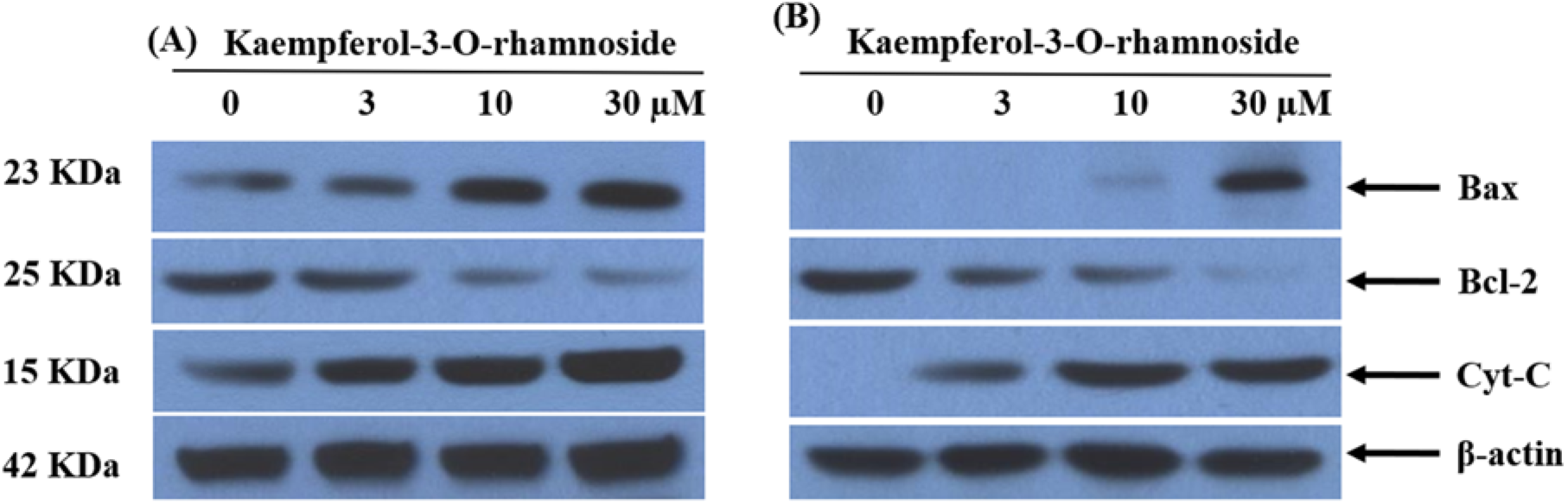
Effect of Kaempferol 3-O-rhamnoside on the expression of Bcl-2, Bax, and Cyt-c in. **A)** HL-60 and **B)** KG-1. Western blotting assays used anti-Bax, Bcl-2, and Cyt-C antibodies; β-Actin was used as an internal control

Western blot analysis using specific antibodies revealed that kaempferol 3-O-rhamnoside, at concentrations ranging from 3 to 30 µM, significantly increased the protein levels of pro-apoptotic Bax and Cyt-c. Conversely, the expression of the anti-apoptotic protein Bcl-2 decreased significantly in a dose-dependent manner, confirming the induction of apoptosis in HL-60 (Fig 5A) and KG-1 cells (Fig 5B). Bax (Bcl-2-associated X protein) is a member of the Bcl-2 family and plays a critical role in regulating apoptosis. Upon activation, Bax translocates from the cytosol to the mitochondrial outer membrane, where it disrupts mitochondrial integrity by forming pores, leading to the release of apoptosis-promoting factors like cytochrome c. This triggers caspase activation and ultimately cell death. Anti-apoptotic proteins such as Bcl-2 and Bcl-xL counteract Bax by inhibiting its activity and stabilizing mitochondrial membrane integrity. The balance between pro-apoptotic proteins (e.g., Bax) and anti-apoptotic proteins (e.g., Bcl-2) is crucial for determining cell fate. Disruption of this balance can contribute to cancer development and progression. Several clinically utilized drugs act by indirectly activating Bax. The Bcl-2 protein family comprises three functional classes, differentiated by the number of Bcl-2 homology (BH) domains they possess: anti-apoptotic proteins (e.g., Bcl-2, Mcl-1, Bcl-xL) with four BH domains, pro-apoptotic effector proteins (e.g., Bax, Bak) also with four BH domains, and “BH3-only” pro-apoptotic proteins (e.g., BID, BIM) with a single BH3 domain. These proteins can form homo- or heterodimers, playing distinct roles in mitochondrial membrane permeability regulation. Bcl-2, specifically, is known for promoting cell survival and growth by preventing apoptosis. Its expression is closely linked to resistance to chemotherapy, as observed in certain lymphomas such as chronic lymphocytic leukemia (CLL), which rely on Bcl-2 for survival. Inhibiting Bcl-2 can facilitate the apoptosis process by allowing cytochrome c release and caspase activation. In this study, Western blot analysis was performed to examine protein expression levels in HL-60 and KG-1 cells treated with kaempferol 3-O-rhamnoside (0–30 μM) for 24 to 48 hours. These results demonstrate that kaempferol 3-O-rhamnoside treatment led to an increased expression of Bax and Cyt-c, as evidenced by darker bands corresponding to Bax in a concentration range of 3–30 μM. Cytochrome c levels also increased gradually over the 24 to 48-hour incubation period. In contrast, Bcl-2 levels decreased significantly, suggesting that kaempferol 3-O-rhamnoside inhibits Bcl-2 expression through protein fragmentation while promoting the formation and activation of Bax and Cyt-c in the mitochondria of HL-60 and KG-1 cells. Several studies have indicated that Bax is a promising direct target for small molecule drug discovery, as direct Bax activators have shown potential in overcoming chemotherapy and radiotherapy resistance. The results of this study highlight kaempferol 3-O-rhamnoside’s ability to modulate the expression of key apoptotic proteins, supporting its potential as an effective agent in cancer therapy targeting apoptosis pathways.

### Molecular docking analysis

PARP1 protein binds to DNA by transferring a chain of ADP-ribosyl residues (PARs) and two zinc finger motifs from nicotinamide-adenine-dinucleotide (NAD+) to chromatin-associated acceptor proteins. This enzyme plays an important role in promoting DNA repair in cancer cells [17]. Therefore, searching for compounds with PARP1 inhibitory activity is considered an effective approach for developing anticancer drugs. In this study, kaempferol-3-O-rhamnoside was found to activate PARP1 in a dose-dependent manner in both HL-60 and KG cells, therefore, it was docked to the catalytic site of PARP1 to evaluate the binding ability and further predict the anticancer mechanism action of this compound (Fig 6).

**Fig 6.**
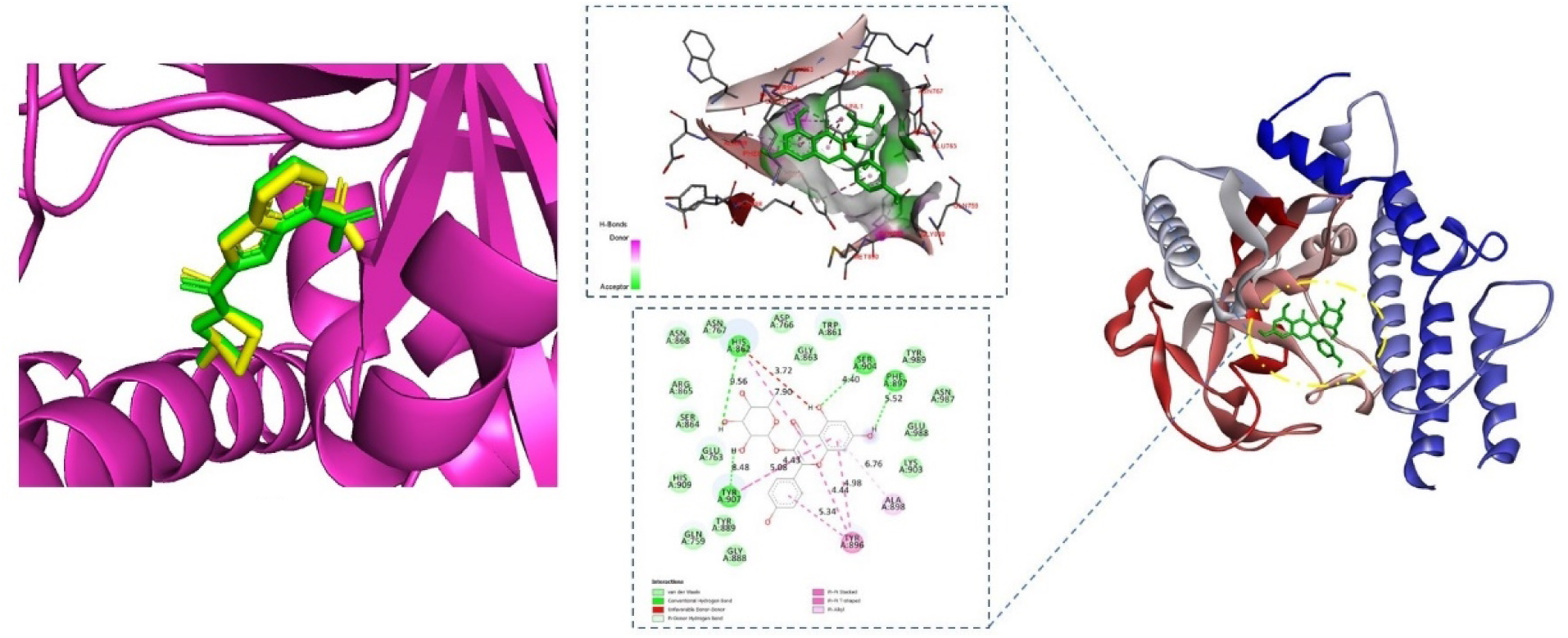
Binding diagram of kaempferol-3-O-rhamnoside with PARP1. Docking pose (yellow) and Crystal pose (green) of reference compound at the crystal structures of PARP1.

The 3D structure of the PARP1 catalytic domain (352 amino acids from THR101 to HIS660) (PDB ID: 5WS1) was selected for screening. The grid box was set to cover the active site of PARP1 according to the position of the co-crystal ligand. The co-crystal ligand, 2-[(3R)-3-azanylpyrrolidin-1-yl] carbonyl-1H-benzimidazole-4-carboxamide, was also docked to validate the selected protein structure and docking by analyzing the RMSD (root mean square deviation) of the docking pose for the crystal pose. As a result, the docking pose had a similar shape and orientation to the crystal pose with a root mean square deviation (RMSD) of 0.525 Å (data not shown). The results showed that kaempferol-3-O-rhamnoside had a docking score of - 10.3 kcal/mol, which was better than the docking score of the reference compound, - 8.6 kcal/mol (Table 2). Kaempferol-3-O-rhamnoside interacted with the enzyme with four hydrogen bonds at HIS862, SER904, PHE897, TYR907, and two hydrophobic interactions at ALA898, TYR896 (Fig 6, Table 2). Overall, the acidic residues of the triad, HIS862 and TYR896, are required for NAD+ binding. While GLU988 plays an important role in catalysis in addition to substrate positioning. Notably, similar to the control compound, kaempferol-3-O-rhamnoside showed interactions with the key amino acid residues of the donor site HIS862 and TYR896 which play an important role in PARP1 activity.

**Table 2.**
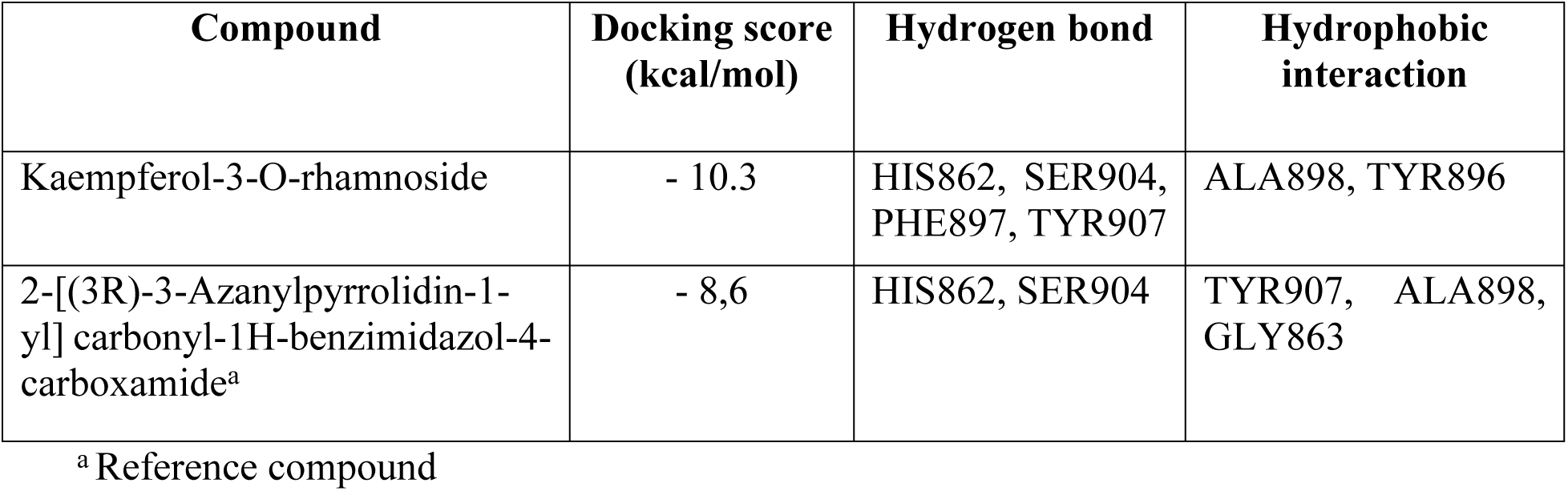
Docking results of testing compounds with PARP1 protein.

### Molecular Dynamics Simulation

Small molecules interacting with protein surfaces can induce significant changes in the tertiary structure, playing a vital role in advancing drug design. One of the key advantages of molecular dynamics (MD) modeling is its ability to assess the flexibility of the protein-ligand complex. This approach offers an assessment of the thermodynamics and kinetics involved in drug-enzyme binding, providing valuable insights into the dynamics of the interaction. In this study, the 100 ns trajectory of the complex between kaempferol-3-O-rhamnoside and PARP1 was analyzed to assess the stability of both the ligand and the protein throughout the simulation (Fig 7).

**Fig 7.**
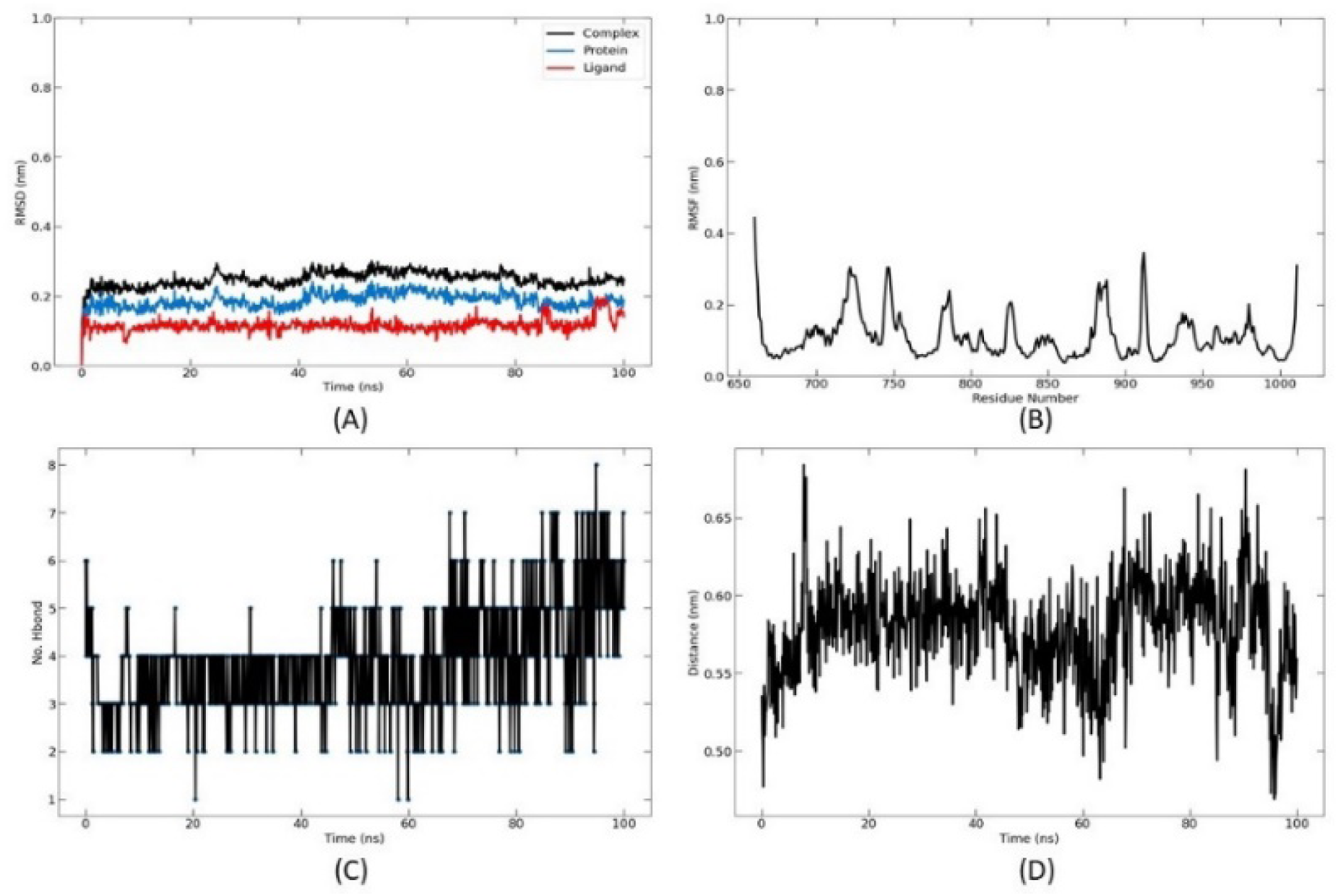
Molecular dynamics simulation of kaempferol-3-O-rhamnoside and PARP1 complex during 100 ns. **A)** RMSD, **B)** RMSF, **C)** Number of hydrogen bonds, and **D)** Average center of mass distance of Kaempferol-3-O-rhamnoside - PARP1 complex.

In RMSD (Root Mean Square Deviation) (Fig 7A), the black line (complex) indicates the overall RMSD of the Kaempferol-3-O-rhamnoside – PARP1 complex. It shows that the system stabilizes after an initial fluctuation phase, suggesting that the complex reaches equilibrium after approximately 20-30 ns. The blue line (protein) shows the RMSD of the PARP1 structure alone, remaining stable throughout, indicating that the protein maintains its structural integrity. The red line (ligand) shows the RMSD of Kaempferol-3-O-rhamnoside, which remains lower compared to the complex and protein, suggesting that the ligand is relatively stable in its binding site without significant shifts. The RMSF plot shows the flexibility of individual amino acid residues throughout the MD simulation (Fig 7B). Peaks in the RMSF graph indicate regions of the protein with higher flexibility, typically loop regions or terminal ends. Lower peaks suggest more rigid or stable regions. Most residues appear to have low RMSF values, indicating minimal fluctuation and a stable binding site. Higher peaks around specific residue numbers may correspond to flexible loops or non-binding regions, which do not significantly affect the stability of the ligand in the active site. About the number of Hydrogen Bonds (Fig 7C), This plot shows the number of hydrogen bonds between kaempferol-3-O-rhamnoside and PARP1 throughout the simulation. The number of hydrogen bonds varies over time, indicating dynamic interactions between the ligand and the protein. The range of 2 to 7 hydrogen bonds during the simulation suggests a moderate to strong interaction, with fluctuations that are common in MD simulations as the ligand and protein adjust and interact under thermal conditions. Center of Mass Distance (Fig 7D) indicates the average distance between the centers of mass of the protein and ligand over time and shows that the distance remains relatively stable around 0.55–0.65 nm, indicating that the ligand remains bound in the binding pocket throughout the simulation. Small fluctuations are typical and indicate minor adjustments in positioning without significant unbinding events. Overall the MD simulation results suggest that kaempferol-3-O-rhamnoside forms a stable complex with PARP1. The low RMSD and consistent hydrogen bond formation indicate good binding stability, the RMSF results show that the binding site is relatively rigid, supporting the ligand’s stable binding. In addition, the stable center of mass distance further confirms that the ligand stays bound throughout the 100 ns simulation. These results imply that kaempferol-3-O-rhamnoside interacts effectively with PARP1, maintaining stability in the binding pocket. This interaction, supported by multiple hydrogen bonds and stable RMSD, suggests potential inhibitory activity and warrants further investigation to confirm its biological relevance.

The radius of gyration (Rg) (Fig 8A) measures the compactness of the protein-ligand complex throughout the MD simulation. The Rg plot shows that the radius remains relatively stable around 2.10–2.15 nm, suggesting that the complex maintains its compact structure throughout the 100 ns simulation. Minimal fluctuation indicates no significant unfolding or loosening of the protein during the binding of kaempferol-3-O-rhamnoside, confirming the structural stability of the complex. Solvent Accessible Surface Area (SASA) (Fig 8B) plot indicates the surface area of the protein-ligand complex accessible to the solvent. The SASA value fluctuates around 175–187.5 nm², showing variability in the exposure of the complex to the solvent. This could be due to minor conformational changes as the protein and ligand adjust during the simulation. These fluctuations are expected in a stable protein-ligand complex and may reflect adjustments to optimize interactions with the solvent. On the Contact Frequency of Residues (Fig 8C), the contact frequency plot shows how often specific amino acid residues contact the ligand during the simulation. Among them, the high Contact Frequency Residues like ASP105, GLU113, PHE108, and SER81 show high contact frequencies, indicating that they are consistently involved in maintaining interactions with the ligand. These residues likely play critical roles in stabilizing the ligand in the binding pocket. There are some moderate Contact Frequency Residues such as THR98, ARG97, HIS862, and TYR907 also show significant contact, suggesting that they contribute to the dynamic interaction network. In addition, some residues show lower contact frequency, indicating transient interactions or peripheral roles in ligand binding. Overall The stable Rg and consistent SASA values support that the kaempferol-3-O-rhamnoside – PARP1 complex remains stable throughout the simulation. The high contact frequency of certain residues as ASP105, PHE108 highlights them as crucial for ligand binding. This information can be used to infer that these residues are part of the binding site and essential for interaction. The consistent contact frequency pattern implies that the ligand is well-positioned within the binding pocket, interacting effectively with key residues throughout the simulation.

**Fig 8.**
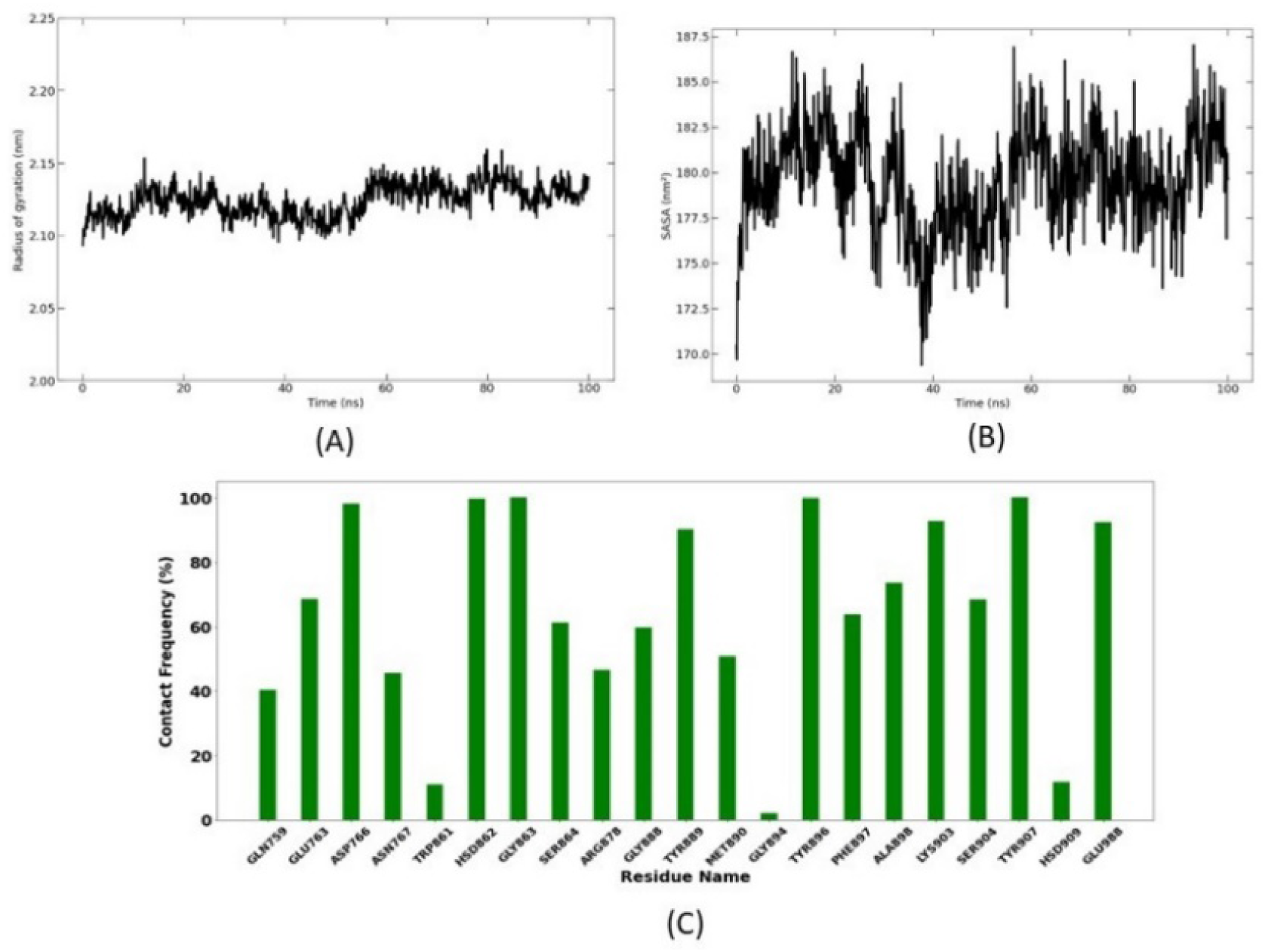
Compactness, folding properties, and contact frequency. **A)** Radius of gyration, **B)** Solvent Accessible Surface Area (SASA), and **C)** Contact histogram of Kaempferol-3-O-rhamnoside - PARP1 complex

In the final stage of the simulation, the binding free energy of kaempferol-3-O-rhamnoside confirmed the results of molecular docking and provided further insight into its inhibitory efficacy against PARP1. The different energy components contributing to the overall binding energy are detailed in Table 3.

**Table 3.** Binding free energy calculation.

| Compounds | Van der Waals energy (kJ/mol) | Electrostatic energy (kJ/mol) | Polar solvation energy (kJ/mol) | SASA energy (kJ/mol) | Binding energy (kJ/mol) |
| --- | --- | --- | --- | --- | --- |
| Kaempferol-3-O-rhamnoside | -185.273 | -164.484 | 235.045 | -20.528 | -135.241 |

In which, Van der Waals Energy (-185.273 kJ/mol) indicates attractive non-covalent interactions between Kaempferol-3-O-rhamnoside and PARP1, suggesting stable interactions within the hydrophobic regions of the binding pocket. This is a crucial indicator of binding potential. Electrostatic Energy (-164.484 kJ/mol) highlights charge attractions between the charged groups of kaempferol-3-O-rhamnoside and the amino acid residues of PARP1. This energy significantly strengthens the binding through interactions such as hydrogen bonding and ionic interactions. Polar Solvation Energy (235.045 kJ/mol) indicates an energy cost associated with the solvation of Kaempferol-3-O-rhamnoside in an aqueous environment, likely due to its polarity and solubility. This energy is counterbalanced by other attractive energy contributions, supporting overall complex stability. The negative solvent-accessible surface area (SASA) energy (-20.528 kJ/mol) suggests that the hydrophobic regions of the ligand are well-protected by the binding pocket, contributing to the stability of the complex in an aqueous medium. Meanwhile, the negative total binding energy (-135.241 kJ/mol) confirms that kaempferol-3-O-rhamnoside has strong binding affinity for PARP1. This energy balance between attractive and repulsive forces supports the stability and binding efficacy of the compound. It is worthy to note that, the significant negative Van der Waals and electrostatic energies, combined with the stabilizing contribution of SASA energy, indicate that kaempferol-3-O-rhamnoside has a strong binding affinity for PARP1. Although there is an energy cost associated with the positive polar solvation energy, the negative total binding energy suggests the stability of the kaempferol-3-O-rhamnoside–PARP1 complex.

## Discussion

Apoptosis is a process of programmed cell death used during early development to eliminate unnecessary cells. In adults, apoptosis plays a crucial role in removing irreparably damaged cells and preventing cancer. When this process is disrupted, uncontrolled cell division may occur, leading to tumor development. Apoptosis is characterized by distinct morphological features, including cell shrinkage, membrane wrinkling, membrane blebbing, and chromatin condensation, which result in the fragmentation of cells into apoptotic bodies [4,5]. In normal conditions, apoptotic bodies are phagocytosed by immune or neighboring cells. This process is regulated by the Bcl-2 family of proteins and cysteine aspartate protease (caspase) enzymes [6–8]. Caspases are divided into two main classes: initiator caspases (caspases 2, 8, 9, 10) and executioner caspases (caspases 3, 6, 7). Initiator caspases generally exist as inactive procaspases, composed of an N-terminal domain and large (p20) and small (p10) subunits [10,14]. The N-terminal domain contains motifs that interact with death-inducing domains of mediator proteins, activating the caspases through oligomerization. This activation triggers the cleavage process, converting executioner caspases from inactive dimers to their active forms. Apoptosis can be initiated via the extracellular pathway through death receptors or the mitochondria-mediated intracellular pathway [7]. The Bcl-2 protein family tightly regulates the activation of the intracellular pathway. Pro-apoptotic proteins such as Bid, Bax, Bim, Bmf, Puma, and Noxa, which contain BH3 domains, promote apoptosis. In contrast, anti-apoptotic proteins like Bcl-2, Bcl-xL, and Mcl-1, which have multiple BH domains, inhibit apoptosis. BH3-only proteins regulate the balance between pro- and anti-apoptotic proteins. Pro-apoptotic proteins increase the permeability of the mitochondrial outer membrane, facilitating the release of cytochrome c, which binds to Apaf-1 and procaspase-9, thereby activating downstream caspases (e.g., caspase-3). Mitochondria also release Smac, a protein that inhibits anti-apoptotic proteins [7,13]. Cancer cell growth and proliferation can be suppressed by regulating the cell cycle and inducing apoptosis, particularly through mitochondrial pathways. Pro-apoptotic proteins such as Bax can form pores in the mitochondrial membrane, allowing cytochrome c to be released into the cytoplasm. Cytochrome c (Cyt-c) then interacts with Apaf-1, activating procaspase-9 to form caspase-9 and assembling the “Apaf-1/caspase-9” holoenzyme complex, which converts procaspase-3 into active caspase-3. This complex maintains caspase-9 in its active form, ensuring optimal protease activity [14,15].

Poly (ADP-ribose) polymerase 1 (PARP1) is an enzyme involved in the repair of DNA single-strand breaks through the base excision repair pathway [16,17,36]. It plays a critical role in maintaining genomic stability by detecting DNA damage and facilitating DNA repair. PARP1 binds to DNA breaks and catalyzes the transfer of ADP-ribose units from NAD+ to target proteins, forming poly(ADP-ribose) chains. This process signals other DNA repair proteins to the site of damage, enabling effective DNA repair and cell survival. In some cases, cancer cells may often experience high levels of DNA damage and genomic instability, and PARP1 aids in repairing this damage, allowing cancer cells to survive and proliferate. Therefore, inhibiting PARP1 in these cells leads to the accumulation of DNA damage, resulting in cell death, a phenomenon known as synthetic lethality [16,17]. Among the PARP1 Inhibitors, Olaparib (Lynparza) is the first PARP1 inhibitor approved for the treatment of BRCA-mutated ovarian and breast cancers. It is used as monotherapy or in combination with other treatments. Others are Rucaparib (Rubraca), Niraparib (Zejula), Talazoparib (Talzenna), and Veliparib [17,18]. Most of the PARP1 inhibitors have some common side effects such as nausea, fatigue, anemia, thrombocytopenia, and gastrointestinal disturbances [19]. Therefore, monitoring and supportive care for managing the side effects of PARP1 inhibitors during cancer treatment are essential. The development and application of PARP1 inhibitors have significantly advanced personalized cancer treatment, offering a promising therapeutic approach for patients with certain types of cancer [19,20].

In this study, we successfully extracted and identified Kaempferol-3-O-rhamnoside, an active ingredient capable of exhibiting cytotoxic effects on leukemia cells such as HL-60 and KG-1 from the medicinal plant *Vối Thuốc* collected in Vietnam. Previously, several studies worldwide have demonstrated the remarkable biological effects of kaempferol-3-O-rhamnoside, including anti-inflammatory, anti-stress, antioxidant, neuroprotective, anti-diabetic, anti-osteoporotic, anti-anxiety, analgesic, and anti-apoptotic activities [32,33]. It has also shown anti-asthma properties by inhibiting the increase of Th2 cytokine (T helper type-2) levels in the lung and bronchoalveolar lavage fluid (BALF) [37]. Additionally, kaempferol-3-O-rhamnoside inhibits cell proliferation in a dose-dependent manner and promotes apoptosis by activating the caspase signaling pathway, including caspase-9, caspase-3, and PARP, in MCF-7 breast cancer cells [38]. In prostate cancer cells, it demonstrated dose-dependent inhibition of proliferation by upregulating the expression of caspase-8, -9, -3, and poly (ADP-ribose) polymerase proteins [39]. Kaempferol-3-O-rhamnoside displayed antiproliferative activity in Jurkat cells with an IC_50_ of 76.3 μM and mildly suppressed IL-2 mRNA expression. Its mechanism of action includes inhibiting Jurkat cell proliferation via Jun Amino-Terminal Kinase signaling [32]. Furthermore, kaempferol-3-O-rhamnoside has shown antimalarial activity by inhibition of Plasmodium falciparum growth [33], efficacy against glucose metabolism disorders in diabetic subjects [40], anti-inflammation, and antitumor [41,42].

To our knowledge, this is the first study to report the antiproliferative and cytotoxic effects of kaempferol-3-O-rhamnoside on leukemia cells. Our *in vitro* results showed that kaempferol-3-O-rhamnoside exhibited significant cytotoxic effects on two human leukemia cell lines HL-60 and KG-1. The compound inhibited cell proliferation in a dose-dependent manner, with notable induction of apoptosis in both cell lines. At molecular levels, this compound activated the caspase signaling cascade, including caspase-9, caspase-3, and cleaved PARP, leading to programmed cell death. This compound increased pro-apoptotic Bax protein levels, decreased anti-apoptotic Bcl-2 protein expression, and induced mitochondrial damage through cytochrome c release, further promoting apoptosis. It is possible to note that, the compound had demonstrated the ability to inhibit cancer cell proliferation and promote apoptosis, particularly in leukemia and other cancer cell lines.

Additionally, computational analysis using *in silico* modeling docking of kaempferol-3-O-rhamnoside on caspase and PARP proteins provides the first predictive insights into its mechanism of action. Investigation of docking indicates that kaempferol-3-O-rhamnoside binds tightly within the active site of PARP1, forming several critical interactions with key amino acid residues via forming hydrogen bonds including ASN107, GLU98, and ASP105. The involvement of hydrophobic amino acid residues such as PHE102 and ILE85 suggests stable hydrophobic interactions that enhance the overall binding affinity. The ligand exhibits a mix of hydrogen bonds, hydrophobic contacts, and possible pi-stacking with PHE108, indicating a stable and multi-faceted interaction within the binding pocket. About the MD Simulation, the RMSD plot shows that the kaempferol-3-O-rhamnoside – PARP1 complex stabilizes after an initial equilibration phase (around 20-30 ns) and remains steady throughout the 100 ns simulation. This suggests that the ligand is well-accommodated within the binding pocket without significant structural deviations.

The RMSF analysis highlights that most residues in the binding site exhibit low fluctuations, indicating a stable environment around the ligand. Peaks corresponding to non-binding regions or flexible loops do not significantly affect the ligand’s stability. The number of hydrogen bonds fluctuates between 2 and 7 over the simulation period, confirming consistent polar interactions that contribute to the stability of the complex. The relatively stable center of mass distance (around 0.55–0.65 nm) between the ligand and the protein suggests that the ligand remains securely in the binding pocket throughout the simulation. The Rg plot shows minor fluctuations around 2.10–2.15 nm, indicating that the complex maintains a compact structure and does not undergo significant conformational changes during the simulation. Moreover, the SASA plot remains fairly consistent, fluctuating around 175–187.5 nm², implying that the ligand’s position within the binding pocket slightly adjusts to optimize interactions with both the protein and solvent. In addition, the energy profile analysis confirms that kaempferol-3-O-rhamnoside interacts strongly and stably with PARP1. The significant contributions from Van der Waals and electrostatic interactions, combined with favorable hydrophobic effects support the inhibitor potential of kaempferol-3-O-rhamnoside to PARP1. This stable binding suggests that kaempferol-3-O-rhamnoside is a promising candidate for further therapeutic exploration.

## Conclusion

Our study reports the isolation of the main active compound from *Schima wallichii* (DC.) Korth and identifies its chemical structure as kaempferol-3-O-rhamnoside. This is the first report of kaempferol-3-O-rhamnoside being isolated from the stem of *Schima wallichii* (DC.) Korth collected in Vietnam. This compound effectively inhibited apoptosis-related growth in HL-60 and KG-1 leukemia cell lines. Specifically, it significantly suppressed cell proliferation and activated caspase-3 and caspase-9 in both HL-60 and KG-1 cells. Kaempferol-3-O-rhamnoside also dose-dependently upregulated the pro-apoptotic protein Bax and downregulated the anti-apoptotic protein Bcl-2 in these cell lines. Docking simulations revealed that Kaempferol-3-O-rhamnoside binds to both the procaspase-3 allosteric site and the active site of PARP1, with binding energies of -7.36 and -10.76 kcal/mol, respectively. In the interaction with PARP1, Kaempferol-3-O-rhamnoside demonstrated stable binding affinity characterized by significant hydrogen bonding, hydrophobic interactions, and pi-stacking. MD simulations confirmed that the complex remained structurally stable throughout the 100 ns period, showing minimal RMSD, Rg, and SASA fluctuations, which indicate the ligand’s robustness in maintaining its binding position. The energy profile further supported these findings, suggesting that binding is driven primarily by Van der Waals and electrostatic forces, balanced by some solvation energy costs. These results imply that Kaempferol-3-O-rhamnoside has potential as a strong PARP1 inhibitor, making it a promising candidate for further *in vitro* and *in vivo* validation. The detailed binding interaction and stability analyses provide a solid foundation for considering this compound in drug discovery and development processes targeting PARP1-related pathways, particularly in cancer therapy. Although it is still early to conclude the leukemia treatment potential of Kaempferol-3-O-rhamnoside from *Schima wallichii*, our finding contributes strong evidence of the cytotoxic potential of *Schima wallichii* against leukemia cells and serves as a basis for future research into the therapeutic applications of this plant.

## Acknowledgments

We would like thank to School of Medicine and Pharmacy and VN-UK Institute for Research and Executive Education, The University of Danang for providing the infrastructure and facilities to perform this study. Furthermore, we sincerely appreciate to Ministry of Education and Training, Vietnam who was providing funds for this research.

## Author Contributions

**Conceptualization:** Manh Hung Tran, Y Nhi Nguyen, Thuy Mi Pham Lam, Thuy Linh Thi Tran, Tan Khanh Nguyen, Tuan Anh Le, Van Ngo Thai Bich, Hieu Phu Chi Truong, Phu Tran Vinh Pham

**Data curation:** Y Nhi Nguyen, Thuy Mi Pham Lam, Thuy Linh Thi Tran.

**Formal analysis:** Thuy Linh Thi Tran, Tan Khanh Nguyen, Tuan Anh Le, Van Ngo Thai Bich, Hieu Phu Chi Truong.

**Funding acquisition:** Manh Hung Tran, Phu Tran Vinh Pham

**Methodology:** Y Nhi Nguyen, Thuy Mi Pham Lam, Thuy Linh Thi Tran.

**Software:** Tan Khanh Nguyen, Tuan Anh Le, Van Ngo Thai Bich, Hieu Phu Chi Truong

**Supervision:** Manh Hung Tran, Phu Tran Vinh Pham.

**Validation:** Manh Hung Tran, Phu Tran Vinh Pham.

**Visualization:** Tan Khanh Nguyen, Tuan Anh Le, Van Ngo Thai Bich, Hieu Phu Chi Truong.

**Writing – original draft:** Y Nhi Nguyen, Thuy Mi Pham Lam, Thuy Linh Thi Tran.

**Writing – review & editing:** Manh Hung Tran, Phu Tran Vinh Pham.

## Funding

This research is supported by the Fund of Ministry of Education and Training, Vietnam under project number B2024.DNA.06

## Competing interests

The authors have declared that no competing interests exist.

## Ethical Approval

Ethical approval is not applicable to this article.

## Statement of Informed Consent

No human subjects are in this article, and informed consent is not applicable.

## Supporting information

**S1 File. NMR spectra of kaempferol-3-O-rhanoside**

## Notes

### Competing Interest Statement

The authors have declared no competing interest.

